# A chimera including a *GROWTH-REGULATING FACTOR* (*GRF*) and its cofactor *GRF-INTERACTING FACTOR* (*GIF*) increases transgenic plant regeneration efficiency

**DOI:** 10.1101/2020.08.23.263905

**Authors:** Juan M. Debernardi, David M. Tricoli, Maria F. Ercoli, Sadiye Hayta, Pamela Ronald, Javier F. Palatnik, Jorge Dubcovsky

## Abstract

Genome editing allows precise DNA manipulation, but its potential is limited in many crops by low regeneration efficiencies and few transformable genotypes. Here, we show that expression of a chimeric protein including wheat GROWTH-REGULATING FACTOR 4 (GRF4) and its cofactor GRF-INTERACTING FACTOR 1 (GIF1) dramatically increases the efficiency and speed of regeneration in wheat, triticale and rice and expands the number of transformable wheat genotypes. Moreover, *GRF4-GIF1* induces efficient wheat regeneration in the absence of exogenous cytokinins, which facilitates selection of transgenic plants without selectable markers. By combining *GRF4-GIF1* and CRISPR-Cas9 technologies, we were able to generate large numbers of edited wheat plants. The *GRF4-GIF1* transgenic plants were fertile and without obvious developmental defects, likely due to post-transcriptional regulatory mechanisms operating on *GRF4* in adult tissues. Finally, we show that a dicot *GRF-GIF* chimera improves regeneration efficiency in citrus suggesting that this strategy can be expanded to dicot crops.

---

Recent studies have reported improvements in plant regeneration efficiency from tissue culture by overexpressing plant developmental regulators including *LEAFY COTYLEDON1* ^1, 2^, *LEAFY COTYLEDON2* ^3^, *WUSCHEL* (*WUS*) ^4^, and *BABY BOOM* (*BBM*) ^5^. Those genes promote the generation of embryo-like structures, somatic embryos or regeneration of shoots. For example, overexpression of the maize developmental regulators *BBM* and *WUS2* produced high transformation frequencies from previously non-transformable maize inbred lines and other monocots species ^6–8^. Another strategy uses different combinations of developmental regulators to induce *de novo* meristems in dicotyledonous species without tissue culture ^9^. Still, there remains a need for new methods providing efficient transformation, increased ease of use, and suitable for a broader range of recalcitrant species and genotypes.

We recently discovered that expression of a sequence encoding a chimeric protein including a GRF transcription factor and its GIF cofactor dramatically increases regeneration efficiency in both monocotyledonous and dicotyledonous species, expands the number of transformable cultivars and results in fertile transgenic plants. *GRF* transcription factors are highly conserved in angiosperms, gymnosperms and moss ^10^. They encode proteins with conserved QLQ and WRC domains that mediate protein-protein and protein-DNA interactions, respectively ^11–13^. Many angiosperm and gymnosperm *GRFs* carry a target site for microRNA miR396, which reduces *GRF*s’ function in mature tissues ^14^.

The GRF proteins form complexes with GIF cofactors that also interact with chromatin remodeling complexes *in vivo* ^15, 16^. Multiple levels of regulation control the efficiency of the assembly of functional GRF/GIF complexes *in vivo* ^17^. Loss-of-function mutations in *GIF* genes mimic the reduced organ size observed in *GRF* loss-of-function mutants or in plants overexpressing miR396 ^11–13, 18, 19^ while overexpression of *GIF* promotes organ growth and can boost the activity of GRFs ^12, 13, 15, 20–22^. Furthermore, simultaneous increases in the expression of Arabidopsis *GRF3* and *GIF1* promotes larger increases of leaf size relative to the individual genes ^15^. Based on the observation that GRFs and GIFs interact to form a protein complex ^15^, we decided to evaluate the effect of a GRF-GIF chimera encoded in a single polypeptide in wheat.

We identified 10 *GRFs* in the wheat genome (Supplementary Figure 1A) and selected wheat *GRF4* based on its homology to *OsGRF4*, a rice gene that promotes grain and plant growth in rice and wheat ^23–27^. Among the three wheat GIF cofactors, we selected the closest homologue of Arabidopsis and rice GIF1 (Supplementary Figure 1B), because members of this clade have been shown to control growth in Arabidopsis, rice and maize ^12, 13, 21, 22^. We then combined *GIF1* and *GRF4* to generate a *GRF4-GIF1* chimera including a short intergenic spacer (Figure 1A) using primers described in Supplementary Table 1 (Supplementary Methods 1). Transgenic plants overexpressing the *GRF4-GIF1* chimera under the maize *UBIQUITIN* promoter (*Ubi::GRF4-GIF1*, Supplementary Method 1) were fertile and showed normal phenotypes (Figure 1B). However, they exhibited a 23.9 % reduction in number of grains per spike and 13.7 % increase in grain weight (Supplementary Table 2).

**Figure 1.**
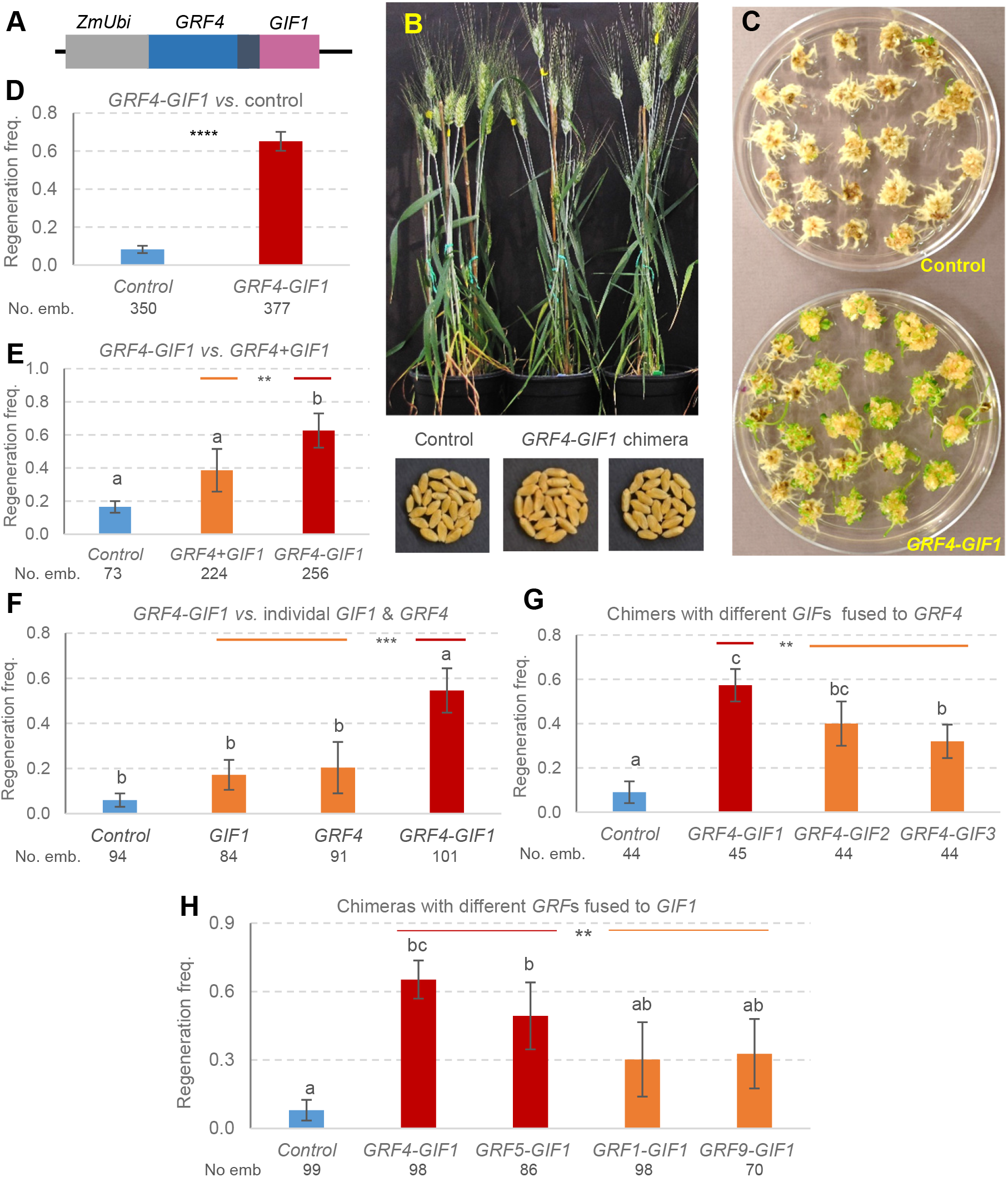
*GRF4-GIF1* chimera. **A)** Schematic representation of the *GRF4* (blue)*-GIF1* (pink) chimera. The black region represents a four amino acid spacer. **B**) The *GRF4-GIF1* transgenic wheat plants were normal and fertile. **C)** Representative transformation showing higher frequency of regenerated shoots during Kronos transformation in the presence of the *GRF4-GIF1* chimera than in the control. **D-H**) Average regeneration frequency of transgenic Kronos plants using experiments as replications and s.e.m. as error bars. All experiments include the empty pLC41 vector as control and the wheat *GRF4-GIF1* chimera. Numbers below the genotypes are total number of inoculated embryos and different letters above bars indicate significant differences (*P* < 0.05, Tukey test). **D**) Control *vs*. *GRF4-GIF1*, n= 14 (**** *P* < 0.0001, square root transformation). **E**) Control, *GRF4-GIF1* and vector including *GRF4* and *GIF1* driven by separate maize *UBIQUITIN* promoters (*GRF4+GIF1*), n = 5 (contrast *GRF4-GIF1 vs. GRF4+GIF1*, ** *P* = 0.0064). **F**) Control, *GRF4-GIF1* and vectors including only *GIF1* or only *GRF4*, n = 5 (contrast *GRF4-GIF1 vs*. combined *GRF4* & *GIF1 P* = 0.0007). **G**) Control and *GRF4* chimeras fused to either *GIF1*, *GIF2* or *GIF3*, n = 3 (contrast chimeras with *GIF1* vs. combined *GIF2* and *GIF3* ** *P* = 0.0046). **H**) Control and chimeras combining different wheat *GRF* genes fused with *GIF1* (n= 4, except for *GRF5* n=3). ** *P* = 0.006 in contrast comparing combined *GRF4-GIF1* and *GRF5-GIF1* chimeras (evolutionary related) with combined *GRF1-GIF1* and *GRF9-GIF1* chimeras (more distantly related). In all tests, normality of residuals was confirmed by Shapiro-Wilk’s test and homogeneity of variances by Levene’s test (raw-data is available in Supplementary Table 3).

We performed 18 transformation experiments in the tetraploid wheat Kronos (Supplementary Methods 2) and estimated regeneration frequencies as the number of calli showing at least one regenerating shoot / total number of inoculated embryos (Supplementary Table 3 summarizes regeneration frequencies and number of inoculated embryos). These regeneration efficiencies were used for five different comparisons using experiments as blocks. Across 15 experiments (Supplementary Table 3), the average regeneration efficiency of the *GRF4-GIF1* chimera (65.1 ± 5.0 %) was 7.8-fold higher than the empty vector control (8.3 ± 1.9 %, *P* < 0.0001, Figure 1C and D).C)

We hypothesize that the increased regeneration efficiency of the *GRF4-GIF1* chimera is associated with the ability of the GRF-GIF complex to regulate the transition between stem cells to transit-amplifying cells ^28^ and their capacity to promote cell proliferation in a broad range of organs ^19^. The wheat *GRF4-GIF1* chimera also accelerates the regeneration process, which allowed us to develop a faster wheat transformation protocol that takes 56 d instead of the 91 d required for all the wheat experiments presented in this manuscript (Supplementary Figure 2).

We then compared the effect on regeneration efficiency of having the *GRF4* and *GIF1* fused in a chimera or expressed separately within the same construct by individual *Ubi* promoters (not fused) (Supplementary Table 3). In five different experiments, the average regeneration efficiency of the separate *GRF4* and *GIF1* genes (38.6 ± 12.9 %) was significantly lower (*P* < 0.0064) than the regeneration efficiency with the *GRF4-GIF1* chimera (62.6 ± 10.3 %, Figure 1E). This result demonstrated that the forced proximity of the two proteins in the chimera increased its ability to induce regeneration.

In another five separate transformation experiments (Supplementary Table 3), we observed significantly lower regeneration efficiencies in embryos transformed with the *GRF4* gene alone (20.4 ± 11.4 %) or the *GIF1* gene alone (17.2 ± 6.6 %) relative to the *GRF4-GIF1* chimera (54.6 ± 9.8 %, contrast *P* = 0.0007, Figure 1F). The regeneration efficiency of the calli transformed with the individual genes was approximately 3-fold higher than the control (6.0 ± 3.0 %) but the differences were not significant in the Tukey test (Figure 1F).

We generated chimeras in which *GIF1* was replaced by other *GIFs* or *GRF4* was replaced by other *GRFs*, and tested their regeneration efficiency in three and four separate experiments, respectively (Supplementary Table 3). The *GRF4-GIF1* combination resulted in higher regeneration efficiency than the *GRF4-GIF2* and *GRF4-GIF3* combination (contrast *P =* 0.0046), and all three chimeras showed higher regeneration efficiency than the control (Tukey test *P* < 0.05, Figure 1G). Similarly, the regeneration efficiency induced by chimeras including the closely related *GRF4* and *GRF5* genes fused with *GIF1*, was higher than the regeneration observed for chimeras including the more distantly related *GRF1* and *GRF9* genes fused with *GIF1* (contrast *P=* 0.0064, Figure 1H). Only the chimeras including the *GRF4* and *GRF5* genes were significantly different from the control (Tukey *P* < 0.05, Figure 1H).

We then tested the potential of the *GRF4-GIF1* chimera to generate transgenic plants from commercial durum, bread wheat and a Triticale line that were recalcitrant to *Agrobacterium*-mediated or had low regeneration efficiency in previous experiments at the UCD Plant Transformation Facility. With the *GRF4-GIF1* chimera we observed high increases in regeneration frequencies in tetraploid wheat Desert King (63.0 ± 17.0 % *vs*. 2.5 ± 2.5 %, 2 experiments) and hexaploid wheat Fielder (61.8 ± 8.2 % *vs*. 12.7 ± 10.3 %, three experiments) relative to the control. For the hexaploid wheat varieties Hahn and Cadenza and the Triticale breeding line UC3190, for which we were not able to generate transgenic plants using the Japan Tobacco protocol, we observed regeneration frequencies of 9 to 19 % with the *GRF4-GIF1* chimera (versus 0 % with the control, Supplementary Figure 3 and Supplementary Table 4A and B).

High wheat regeneration efficiencies have been reported before using the proprietary Japan Tobacco method in the variety Fielder ^29, 30, 31^. However, the company warns that these high values require the optimization of multiple factors with narrow optimal windows and that “those values can drop drastically when one of the factors become suboptimal” ^29^ (Supplementary Table 5). The addition of the *GRF4-GIF1* chimera overcame some of the constrains imposed by these narrow optimal windows and allowed us to obtain high transformation efficiencies using a shorter protocol and embryos of a wider range of sizes (1.5 to 3.0 mm) obtained from plants grown in diverse environmental conditions. High regeneration efficiencies were observed even when we used different vectors and genotypes and without embryo excision, a critical step in the Japan Tobacco technology ^29^.

To test the robustness of our method, we transferred our *GRF4-GIF1* vector to the John Innes Centre Transformation facility for testing with their recently published wheat transformation method ^32^. Fielder plants transformed with the *GRF4-GIF1* chimera showed a 77.5% regeneration efficiency, compared with 33.3% in the control (Supplementary Table 4A). Taken together, these results indicate that the addition of the *GRF4-GIF1* chimera increases the robustness of wheat transformation under different conditions and protocols.

We also tested the wheat *GRF4-GIF1* chimera in the rice variety Kitaake (Supplementary Methods 3). In four independent transformation experiments, we observed a 2.1-fold increase in rice regeneration efficiency (*P* < 0.00001) in the calli transformed with the wheat *GRF4-GIF1* chimera (average 42.8 ± 2.6 %) compared with those transformed with the control vectors (20.3 ± 2.9 %, Supplementary Table 6). These results suggest that the wheat *GRF4-GIF1* chimera is effective in enhancing regeneration in another agronomically important monocotyledonous species.

In many plant transformation systems cytokinins are required to regenerate shoots (Figure 2A). Interestingly, in both laboratories we observed that Kronos and Fielder embryos inoculated with *Agrobacterium* transformed with the *GRF4-GIF1* chimera were able to rapidly regenerate green shoots in auxin media without cytokinin (Figure 2B). We then tested the regeneration efficiency of immature embryos from stable *GRF4-GIF1* transgenics (n=27) and non-transgenic (n=26) T_1_ sister lines in the absence of cytokinin and hygromycin. Under these conditions, the regeneration efficiency of the *GRF4-GIF1* transgenic plants (77.8 %) was significantly higher than the non-transgenic sister lines (11.5 %, Supplementary Figure 4). These results indicated that the *GRF4-GIF1* chimera can promote either embryogenesis, shoot proliferation, or both, in wheat without the addition of exogenous cytokinin.

**Figure 2.**
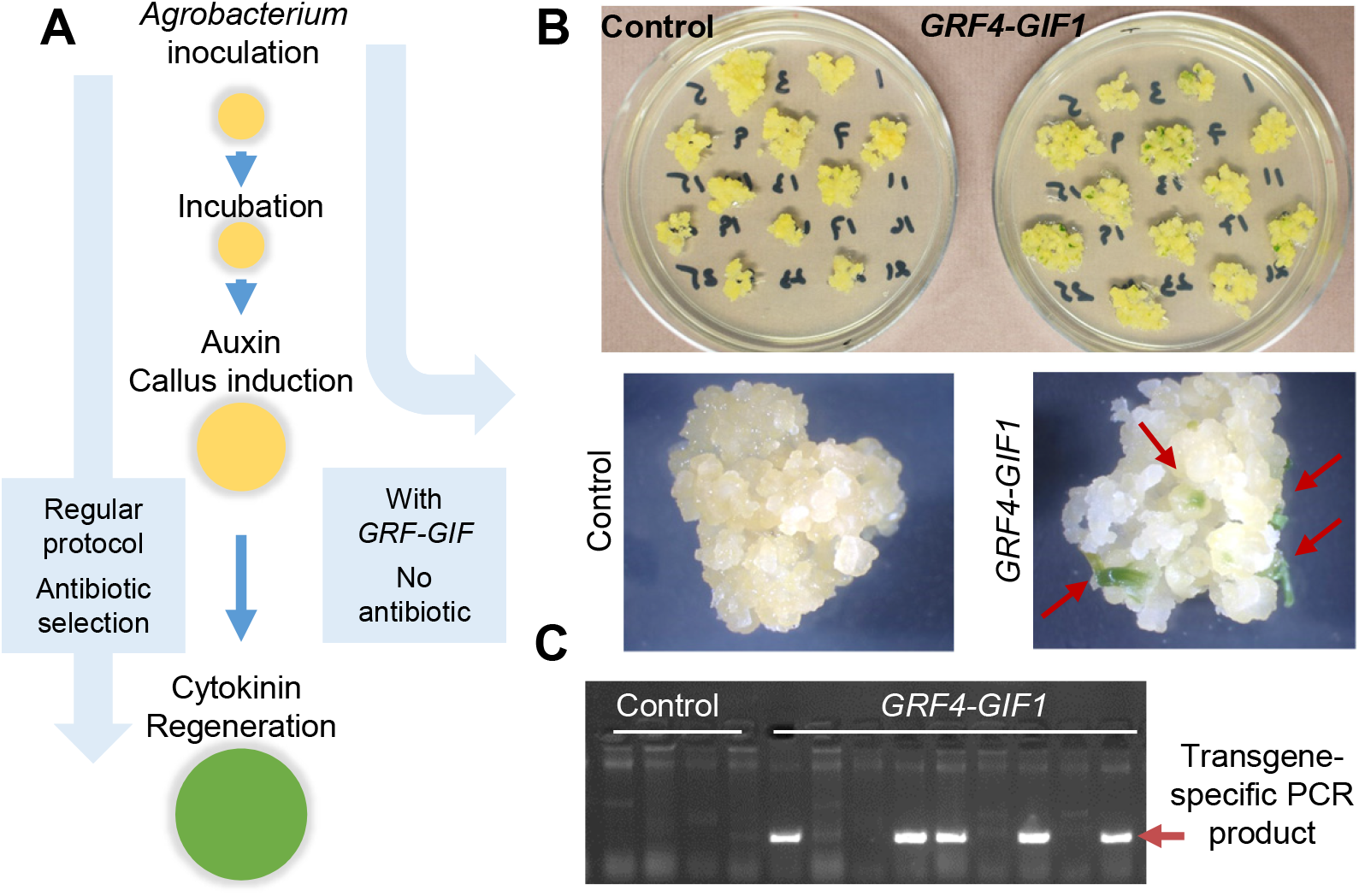
The *GRF4-GIF1* chimera induces embryogenesis in the absence of cytokinins. **A)** Schematic representation of the different steps of wheat transformation. **B)**. Representative calli in auxin media with no hygromycin. Note growing green shoots in callus transformed with the wheat *GRF4-GIF1* chimera in the absence of cytokinins (red arrows). Control: pLC41. **C)** Transgenic specific PCR product (arrow) shows no transgenic plants among four plants regenerated from the control and five transgenic plants among nine regenerated from the *GRF4-GIF1* chimera.

Based on the previous result, we developed a protocol to select transgenic shoots in auxin media without using antibiotic-based markers. In three experiment we recovered 40 shoots using a *GRF4-GIF1* marker-free vector and 15 for the empty vector. Genotyping revealed that 10 out of the 40 (25 %) *GRF4-GIF1* shoots were transgenic, while none of the control was positive (Figure 2C). These high-regenerating transgenic plants overexpressing the *GRF4-GIF1* chimera without selection markers could potentially be used for future transformation experiments to incorporate other genes using selectable markers. This approach could generate separate insertion sites for the *GRF4-GIF1* and the second transgene, facilitating the segregation of the *GRF4-GIF1* insertion in the next generation.

This strategy is not necessary for genome editing, since both the CRISPR-Cas9 and *GRF4-GIF1* sequences can be segregated out together after editing the desired region of the genome. Therefore, the GRF-GIF system is ideal to expand the utilization of genome editing technology to crops with low regeneration efficiencies. As a proof of concept, we generated a binary vector for *Agrobacterium* transformation that contained a cassette including the *GRF4-GIF1* chimera, Cas9 and a gRNA targeting the wheat gene Q (= *AP2L5*) ^33^ in the same T-DNA region (Figure 3A and B). We recovered 30 independent transgenic events out of 32 infected calli (93.7% efficiency, Figure 3C). Disruption of a *Sty*I restriction sites showed Cas9-induced editing in all 30 transgenics (Supplementary Figure 5). We sequenced the PCR products obtained from 10 independent lines and confirmed editing (Figure 3D). Of the ten edited T_0_ plants transferred to soil seven showed clear mutant *q*-null phenotypes (Figure 3E) and the other 3 died before heading. These T_0_ transgenic plants showed normal fertility and the edited *Q* gene and the CRISPR-Cas9 / *GRF4-GIF1* construct are expected to segregate in the T_1_ progeny, facilitating the selection of edited plants without the transgene.

**Figure 3.**
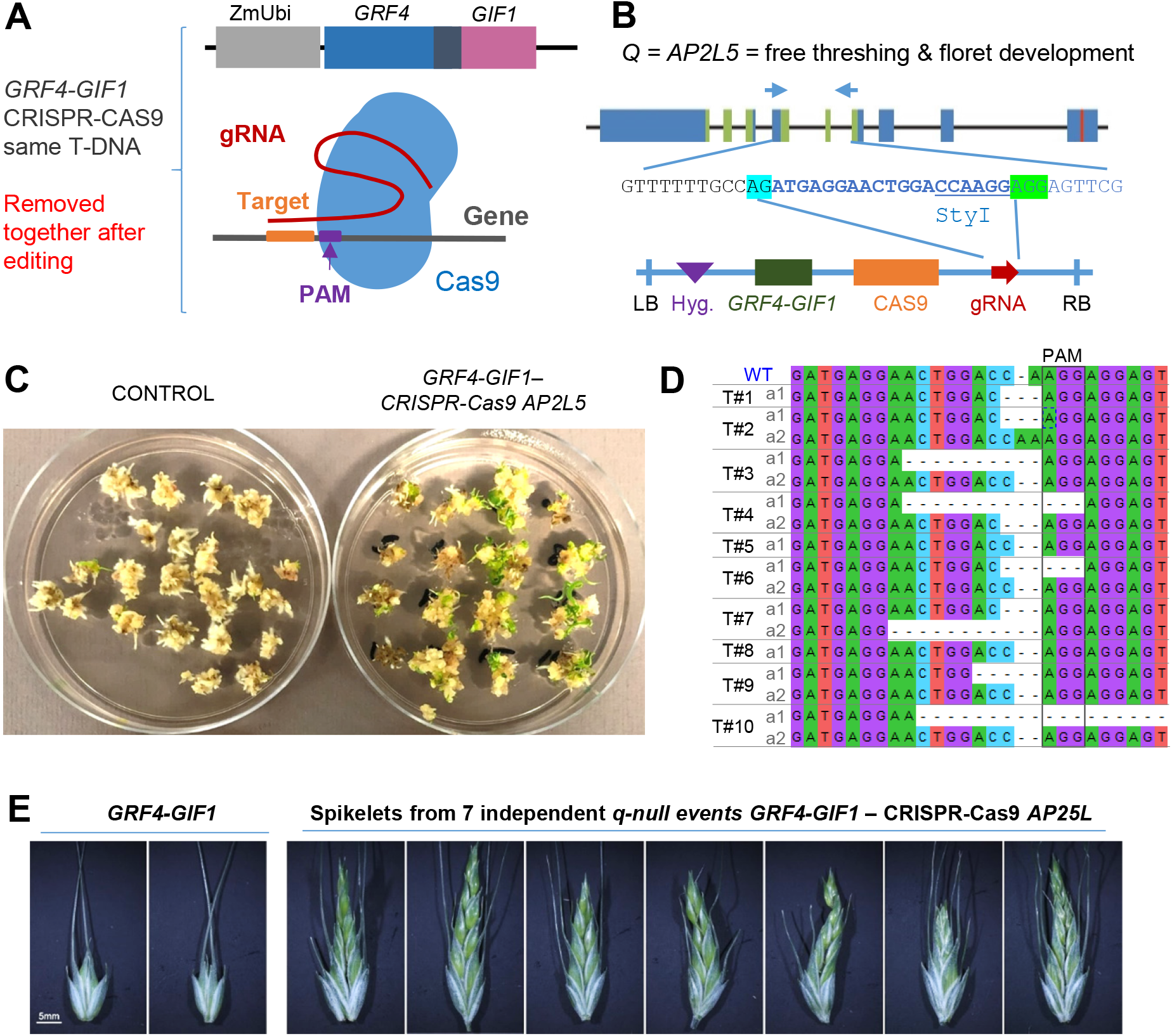
High frequency of genome edited plants using combined GRF4-GIF1 – CRISPR-Cas9 technology. **A)** Technologies combined in a single vector. **B)** Region of the gene *Q* (*AP2L-A5*) targeted with the guide RNA (gRNA) and schematic representation of the vector combining both technologies (LB = left border, Hyg. = hygromycin resistance, RB = right border). **C)** Kronos shoot regeneration of embryos transformed with an empty vector and with the combined GRF4-GIF1 - CRISPR-Cas9-gRNA-*AP2L-A5* construct (93.7 % regeneration efficiency). **D)** All 10 sequenced transgenic T0 plants showed *AP2L-A5* editing. Seven of the 10 plants (T#1 to T#10) carried two different mutations (a1 and a2), documenting high editing efficiency. **E**) Edited T0 plants showed increased number of florets per spikelet (characteristic of *q*-null plants).

Lastly, we performed a series of *Citrus* transformation experiments to test the effect of the *GRF-GIF* technology in a dicot crop with limited regeneration efficiency and organogenic-based transformation protocols. We generated a citrus and a heterologous grape *GRF-GIF* chimera using the closest homologs to wheat *GRF4* and *GIF1* in both species (Supplementary Figure 1A and B). In three independent transformation experiments in the citron rootstock Carrizo (Supplementary Methods 4), epicotyls were transformed with the citrus and the grape *GRF-GIF* chimeras. Epicotyls transformed with the citrus *GRF-GIF* chimera showed a 4.7-fold increase in regeneration frequency relative to those transformed with the empty vector control (Supplementary Figure 6A). The heterologous grape *GRF-GIF* chimera produced similar increases in citrus regeneration efficiency as the citrus chimera (Supplementary Figure 6B).

We also tested the effect of a miR396-resistant grape *GRF-GIF* version (henceforth, *rGRF-GIF*), in which we introduced silent mutations in the *GRF* binding site for miR396 to avoid cleavage (Supplementary Figure 6B and C). In three independent experiments, we observed that the grape *rGRF-GIF* chimera produced the highest frequency of transgenic citrus events (7.4-fold increase compared to the control, *P* < 0.05). A statistical analysis comparing the control versus the three combined GRF-GIF constructs was also significant (*P* = 0.0136, Supplementary Figure 6D and Supplementary Table 7). In spite of its higher-regeneration frequency, the *rGRF-GIF* construct would require additional optimization (e.g. an inducible system) because some of the transgenic events produced large calli that were unable to generate shoots (Supplementary Figure 6B).

In summary, the expression of a *GRF4-GIF1* chimera increased significantly the efficiency and speed of wheat regeneration and the ability to generate large numbers of fertile edited plants, expanded the range of transformable genotypes and eliminated the requirement of cytokinin for regeneration, thereby eliminating the need of antibiotic-based selectable markers. The *GRF4-GIF1* technology results in fertile and normal transgenic plants without the need of specialized promoters or transgene excision, overcoming some of the limitations of transformation technologies with other morphogenic genes (Supplementary Table 8). Because *GRF4-GIF1* likely operates at a later stage of meristem differentiation and stem cell proliferation ^28^ than *Bbm-Wus2* ^6–8^, there is potential to combine both technologies and have synergistic effects in the regeneration efficiency of recalcitrant genotypes. A concurrent and independent work showed that overexpression of Arabidopsis *AtGRF5* and *AtGRF5* homologs positively enhance regeneration and transformation in monocot and dicot species not tested here ^34^. We hypothesize that the benefits of the *GRF4-GIF1* technology can be rapidly expanded to other crops with low regeneration efficiencies by incorporating the *GRF4-GIF1* chimera into currently available protocols. This hypothesis is supported by the high conservation of the GRF and GIF proteins across the plant kingdom and by the enhanced regeneration frequency observed for rice and citrus in this study.

## Online content

Supplementary methods, figures and tables are available in the Supplementary Materials.

## Acknowledgements

This project was supported by the Howard Hughes Medical Institute, NRI Competitive Grant 2017-67007-25939 from the USDA National Institute of Food and Agriculture (NIFA) and the International Wheat Partnership Initiative (IWYP). J.F.P. acknowledges support from the Argentinean Research Council (CONICET) and Agencia Nacional de Promoción de la Investigación, el Desarrollo Tecnológico y la Innovación. P.C.R. was supported by NIH grant #GM122968. S.H. acknowledges support from the Biotechnology and Biological Sciences Research Council Genes in the Environment Institute Strategic Programme BB/P013511/1. J.M.D. was supported by a fellowship (LT000590/2014-L) of the Human Frontier Science Program. M.F.E. is a Latin American Fellow in the Biomedical Sciences, supported by the Pew Charitable Trusts. We thank Dr. Yanpeng Wang for the pYP25F binary vector and Mariana Padilla, Gonzalo Rabasa, Bailey Van Bockern, and Mark Smedley for excellent technical support, and to Cristobal Uauy for coordinating the testing of the *GRF4-GIF1* chimera at the John Innes Centre.

## Author Contributions

**Juan M. Debernardi**: Investigation, Methodology, Formal analysis, Writing - Original Draft - Review & Editing. **David M. Tricoli**: Investigation, Supervision, Methodology, Project administration and funding acquisition, Writing - Review & Editing. **Javier F. Palatnik**: Conceptualization, Writing - Review & Editing. **Maria F. Ercoli**: Investigation (rice section). Writing - Review & Editing. **Sadiye Hayta**: Investigation (JIC wheat transformation). **Pam Ronald**: Supervision (rice section), Writing - Review & Editing. **Jorge Dubcovsky**: Conceptualization, Formal analysis, Supervision, Project administration and funding acquisition, Writing - Original Draft -Review & Editing.

## Data availability statement

Accession numbers and gene names are available in the phylogenetic tree in Supplementary Figure 1. All wheat gene names are based on genome release RefSeq v1.0. The raw data for the different experiments is available in Supplementary Tables 3-4 and 6-7. The methods for the generation of the different vectors and the transformation protocols are described in Supplementary Methods.

