## Supplementary Tables and Figures for "A chimera including a *GROWTH-REGULATING FACTOR* (*GRF*) and its cofactor *GRF-INTERACTING FACTOR* (*GIF*) increases transgenic plant regeneration efficiency"

#### Supplementary Figures

**Supplementary Figure 1.** Phylogenetic trees of GRF and GIF families for wheat (yellow highlight and corresponding RefSeq v1.0 names), rice, Arabidopsis, citrus and grape. The closest homologs to wheat GRF4 and GIF1 are highlighted in orange for citrus and in violet for grape. **A**) We used the QLQ and WRC domains for the analysis of the GRF proteins and **B**) the SNH domain for the analysis of the GIF proteins. The evolutionary history was inferred by using the Maximum Likelihood method. We show the tree with the highest log-likelihood. The percentage of trees in which the associated taxa clustered together is shown next to the branches. We conducted the evolutionary analysis in MEGA X [1]. Yellow highlight: wheat. Orange highlight: selected *Citrus* homolog. Violet highlight: selected *Vitis* homolog. Note that the cluster including wheat and rice GRF3, GRF4, and GRF5 proteins does not include any Arabidopsis protein.

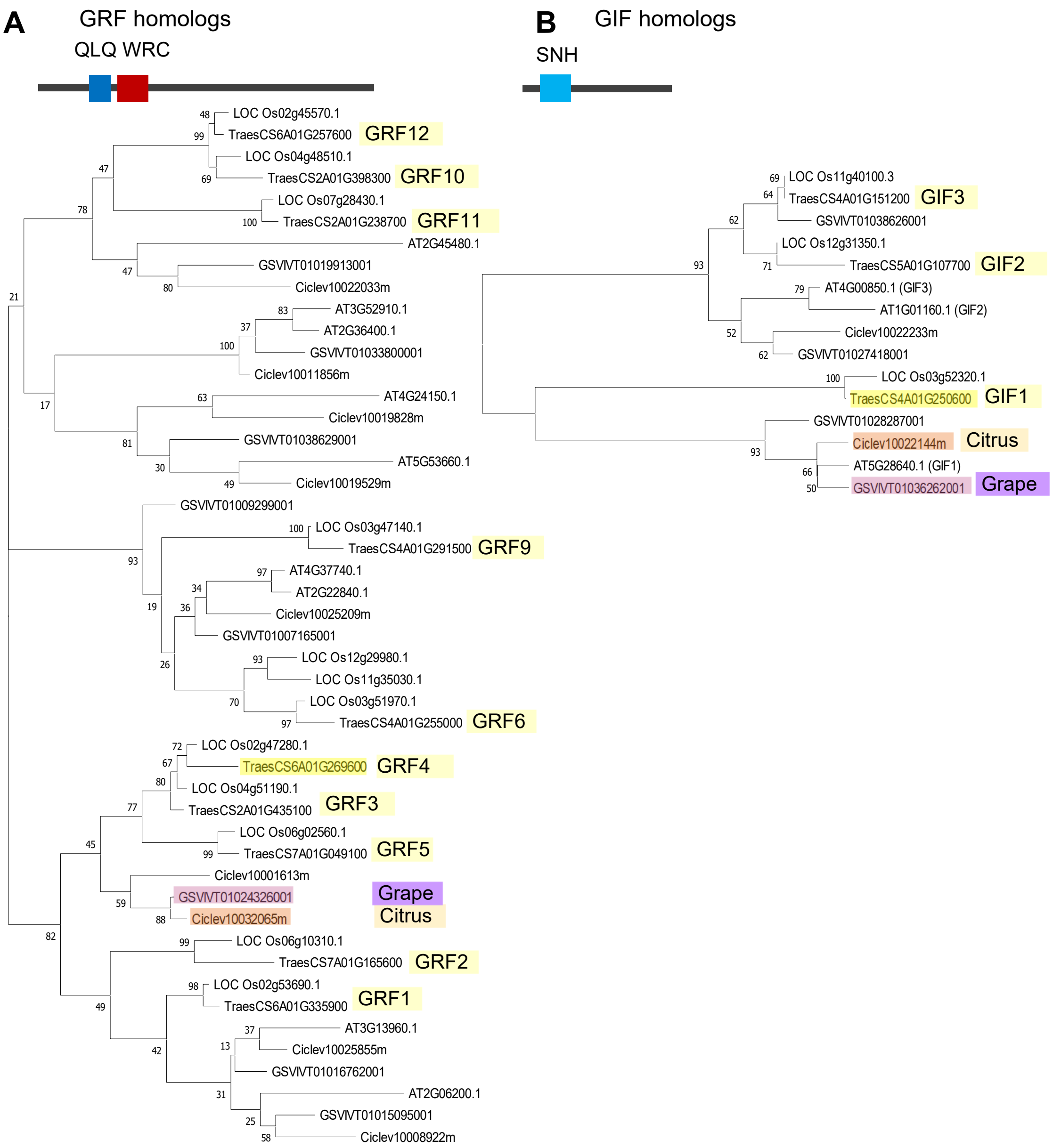

**Supplementary Figure 2.** Accelerated wheat transformation protocol using the *GRF4-GIF1* chimera relative to normal protocol of wheat transformation at the UC Davis transformation facility. The protocol with the *GRF4-GIF1* chimera is faster, reducing the overall process by 5 weeks.

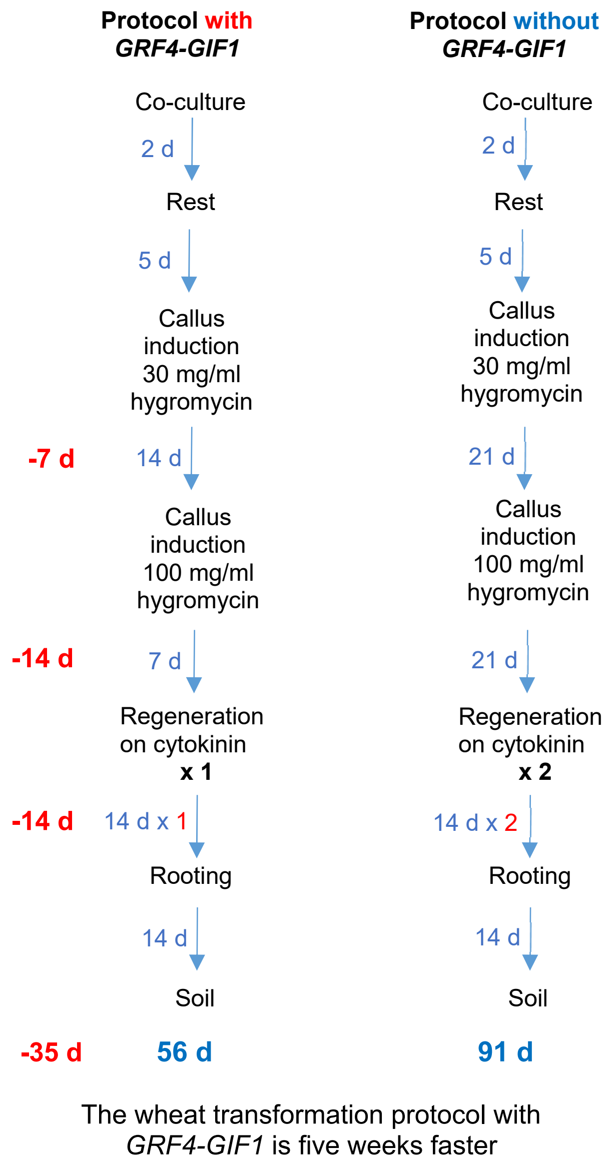

**Supplementary Figure 3.** Effect of the *GRF4-GIF1* chimera in regeneration efficiency in different genotypes. **A**) Representative transformations showing higher frequency of regenerated shoots in the presence of the *GRF4-GIF1* chimera than in the control (empty vector) in different wheat and Triticale genotypes. **B**) Regeneration efficiencies of *GRF4-GIF1 vs.* control in the same cultivars as in A. The raw data is available in Supplementary Table 4A and B. The number of independent experiments is indicated in parenthesis after the genotype name and the total number of inoculated embryos is indicated below. Error bars are s.e.m. No statistical analysis is presented for these experiments because transformations of these cultivars without the *GRF4-GIF1* chimera showed 0 or close to 0 regeneration frequencies.

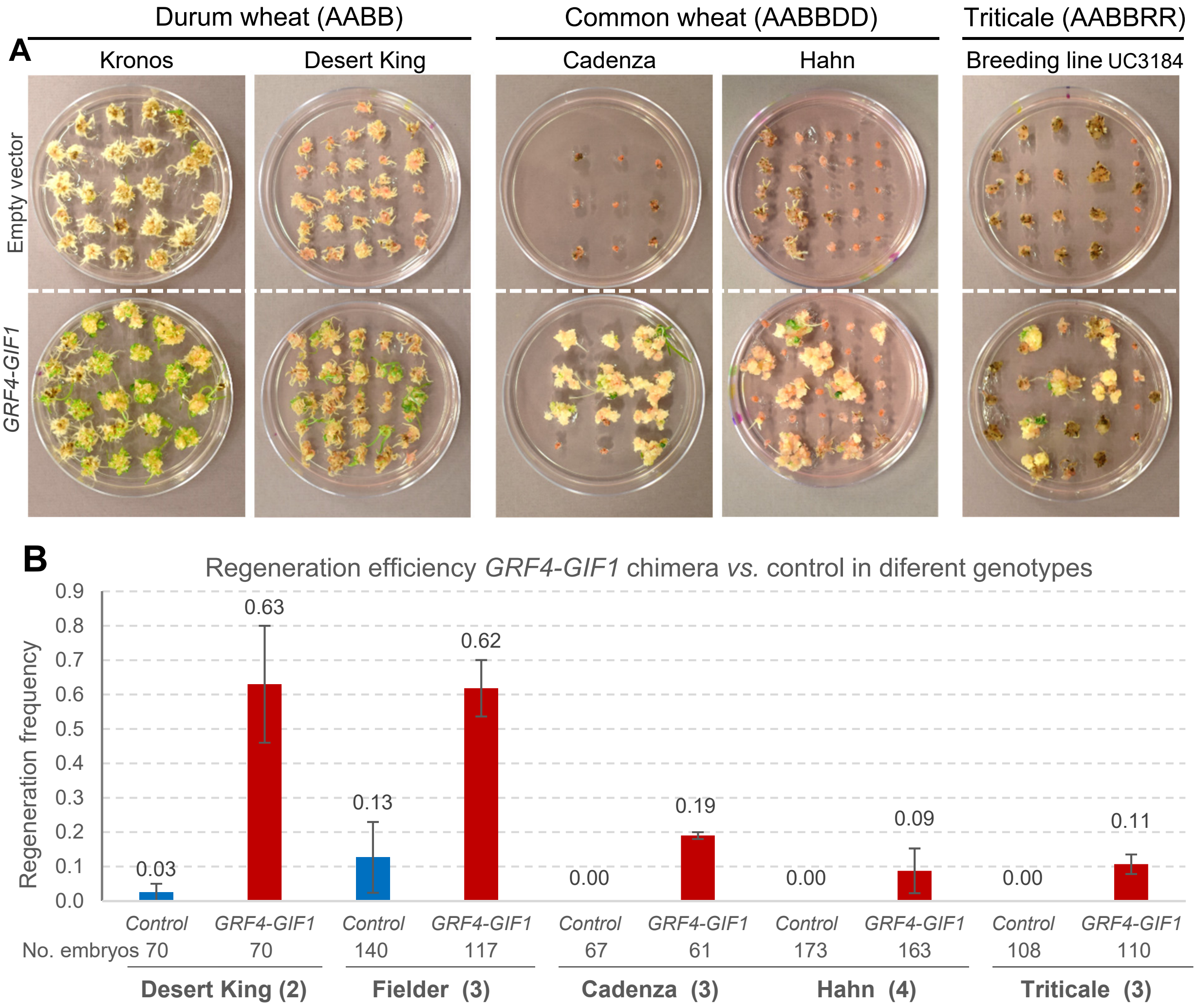

**Supplementary Figure 4.** Effect of the *GRF4-GIF1* chimera in regeneration efficiency in the absence of exogenous cytokinin. Immature wheat embryos from a *GRF4-GIF1* transgenic Kronos $T_1$ plant and a segregating non-transgenic $T_1$ sister line where treated following the standard transformation protocol, excluding the *Agrobacterium* inoculation and the addition of hygromycin to the plates. In the last step, the calli where transferred to regeneration media in the absence of cytokinin. The number of calli regenerating green shoots was significantly higher in the *GRF4-GIF1* transgenic plant (21 out of 27) than in the non-transgenic sister control (3 out of 26). The picture shows representative plates with calli in regeneration media without cytokinin.

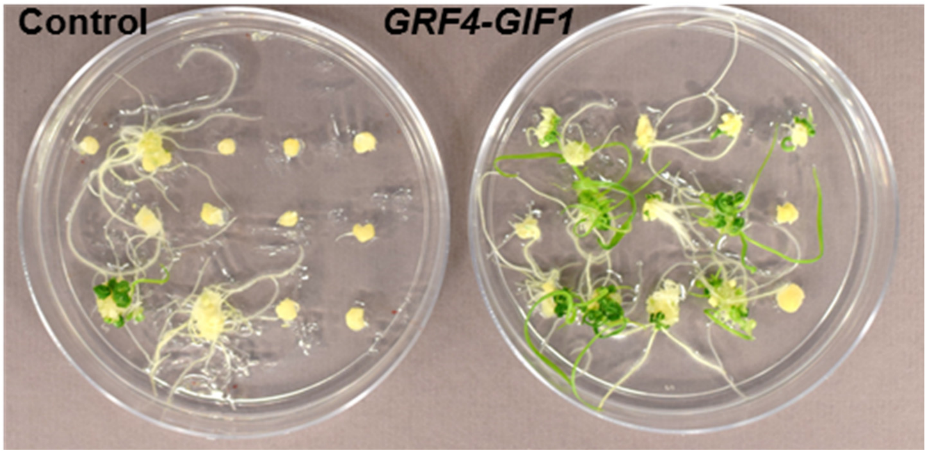

**Supplementary Figure 5. Genome edited wheat plants using combined *GRF4-GIF1* – CRISPR-Cas9 technology**. We recovered 30 independent transgenic events (white numbers) out of 32 infected callus. Calli 6 and 13 were not transgenic. Figures are examples of a transgenic (2) and non-transgenic (13) callus. Editing disrupts a *Sty*I restriction site in the target region resulting in an undigested band (red arrow). Not edited sequences are digested (blue arrows). Ten of the edited events were sequenced and the detected mutations are presented in Figure 3D.

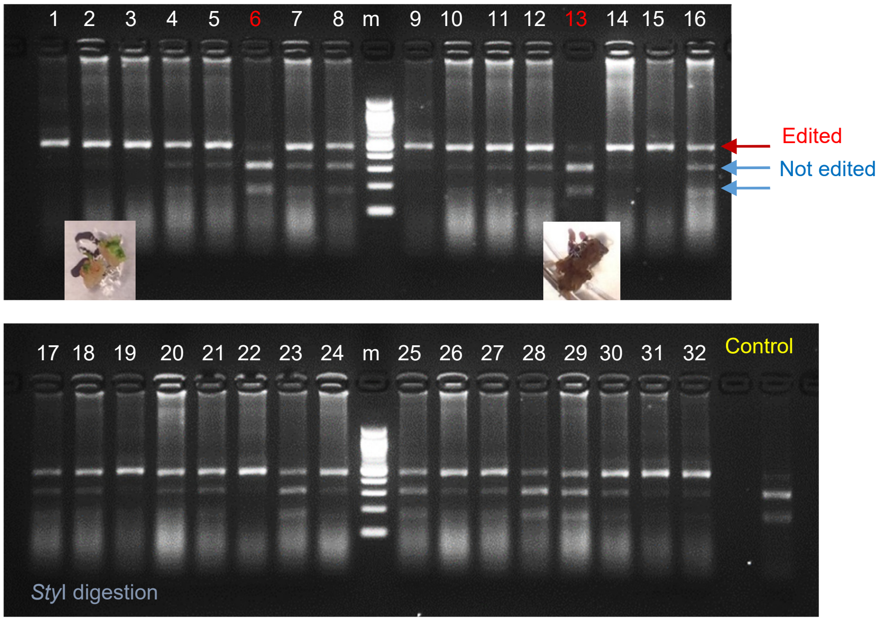

**Supplementary Figure 6. Transformation of dicot species with GRF-GIF chimers. A**) *Citrus* epicotyls transformed with an empty vector and the *Citrus GRF-GIF* chimera (60 d after *Agrobacterium* inoculation). **B**) *Citrus* epicotyls transformed with an empty vector and the *Vitis GRF-GIF* and miR396-resistant *Vitis GRF-GIF* (*rGRF-GIF*) (120 d after inoculation). **C**) Scheme of a *Vitis GRF-GIF* chimera showing the miR396 target site and its interaction with miR396 below. In the miR396-resistant *rGRF-GIF* version, we introduced silent mutations (in red) to reduce interactions with miR396. **D**) Statistical comparison of the three *Citrus* experiments. Different letters above the bars indicate significant Tukey test ($P < 0.05$). Error bars are s.e.m. Horizontal lines on top indicate a significant contrast between the control and combined *GRF-GIF* constructs ($P = 0.0136$). The number of independent experiments is indicated in parenthesis after names and the number of inoculated epicotyls below. Normality of residuals was confirmed by Shapiro-Wilk's test and homogeneity of variances by Levene's test.

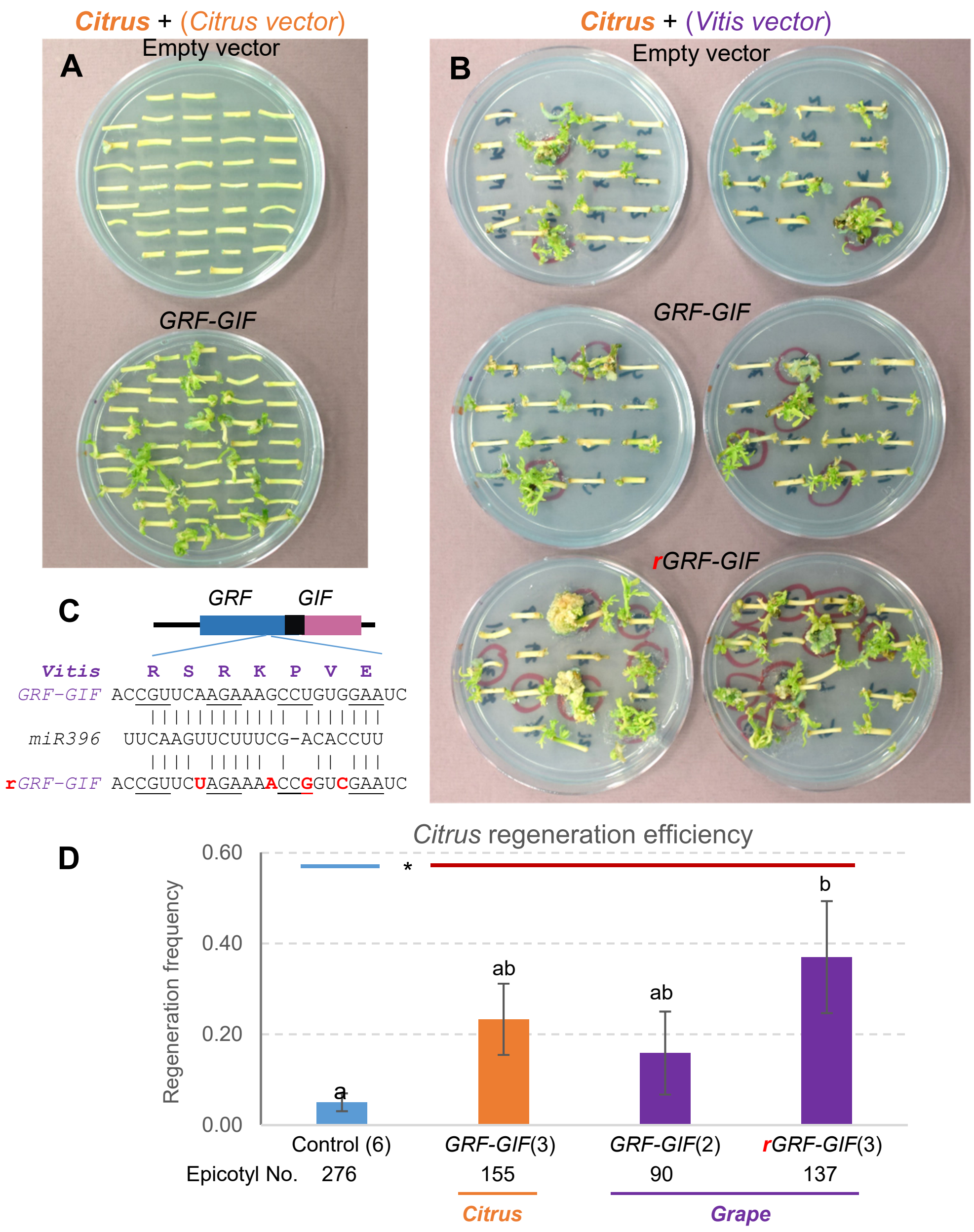

#### SUPPLEMENTARY TABLES

**Supplementary Table 1**. Primers used in this study.

| Name | Sequence | Gene |
|---|---|---|
| Fw-GRF4a | GGGGACAAGTTTGTACAAAAAAGCTGCCACCATGGCGATGCCGTATGCCTCT | GRF4 |
| Rev-GRF4a | GGGGACCACTTTGTACAAGAAAGCTGAACGGTACATYTCGCCGGCGAACAG | |
| Fw-GIF1a | GGGGACAAGTTTGTACAAAAAAGCTGCCACCATGCAGCAGCAACACCTGATG | GIF1 |
| Rev-GIF1a | GGGGACCACTTTGTACAAGAAAGCTGAACGGCTTCCTTCCTCCTCGGT | |
| Fw-GRF4a | GGGGACAAGTTTGTACAAAAAAGCTGCCACCATGGCGATGCCGTATGCCTCT | GRF4-GIF1 |
| Rev-GRF4b | GGCAGCGGCCGCGTACATYTCGCCGGCGAACAG | |
| Fw-GIF1b | GCGGCCGCTGCCATGCAGCAGCAACACCTGATG | |
| Rev-GIF1b | GGGGACCACTTTGTACAAGAAAGCTGAACGCTAGCTTCCTTCCTCCTCGGT | |
| Fw_HindIII | GCCACTCAGCAAGCTTTGCAGCGT | Ubi::GRF4-term |
| Rev-HindIII | TCACGCTGCAAAGCTCTAATTCCCGATCTAGTAAC | |
| Fw_ZmUbi-AscI | GGATCTGCAGGCGCGTGCAGCGTGACCCGGTCGTG | Ubi::GRF4-GIF-term |
| Rev_NosTerm-AscI | TGCACTGCAGGCGCGCTAATTCCCGATCTAGTAAC | |
| QT1-F-GG | ACTTGATGAGGAACTGGACCAAGG | Q gene gRNA |
| QT1-R-GG | AAACCCTTGGTCCAGTTCCTCATC | |
| QT1check-F | TGAGCGACTACGAGGAGGAT | Q gene genotyping |
| QT1check-R | CAGCTGCCCTGTCACATCTA | |
| Fw-rGRF-Vvi | TCTAGAAAACCGGTCGAATCACAAACTA | rGRF-GIF |
| Rev-rGRF-Vvi | TCGACCGGTTTTCTAGAACGGTTGCGG | |
| Fw-GRF-Vvi | GGGGACAAGTTTGTACAAAAAAGCTGCCACCATGAAGCAAAGCTTTGTGG | |
| Rev-GIF-Vvi | GGGGACCACTTTGTACAAGAAAGCTGAACGTCAATTCCCATCTTCAGCA | |
| pLC41_1064 | TCGCTTATTTAAAGGGCGAAT | Transgenic plants genotyping |
| pLC41_1061 | AGCGCGCAAACTAGGATAAA | |

**Supplementary Table 2.** Grain measurements in *GRF4-GIF1* T1 transgenic plants and their sister negative controls in a growth chamber (16 h light at 22 °C and 8 h darkness at 18 °C, light intensity 260 µM $m^{-2}$ $s^{-1}$). Two statistical analyses are presented: 1) A more conservative test using the averages of the families from each event as experimental units (5 negatives *vs*. 3 positives). 2) A more liberal test using the individual plants as experimental units (38 negatives *vs*. 16 positives). Parameters from individual plants were obtained from an average of 23 grains estimated from a Marvin Grain Analyzer. The JD561 numbers indicate independent transformation events with the same *GRF4-GIF1* construct.

| *GRF4-GIF1* (JD561) | Plants | Spikelets / spike | Grains / spike | TGW(g) | Area ($mm^2$) | Width (mm) | Length (mm) |
|---|---|---|---|---|---|---|---|
| Negative (JD561#2-1) | 8 | 13.25 | 26 | 41.48 | 17.75 | 3.31 | 7.43 |
| Negative (JD561#12-1) | 9 | 12.78 | 23 | 57.04 | 20.85 | 3.69 | 7.80 |
| Negative (JD561#20-6) | 4 | 11.50 | 23 | 55.92 | 20.07 | 3.66 | 7.63 |
| Negative (JD561#21-1) | 12 | 11.67 | 27 | 55.50 | 19.74 | 3.60 | 7.63 |
| Negative (JD561#23-8 ) | 5 | 12.80 | 26 | 51.81 | 19.27 | 3.55 | 7.43 |
| Positive (JD561#13-1) | 9 | 9.67 | 18 | 59.68 | 22.18 | 3.67 | 8.31 |
| Positive (JD561#20-11) | 2 | 10.00 | 13 | 57.09 | 21.60 | 3.69 | 8.08 |
| Positive (JD561#23-6) | 5 | 11.00 | 21 | 60.66 | 21.38 | 3.68 | 8.00 |
| Weighted Avg. negatives | 38 | 12.39 | 25.10 | 52.47 | 19.56 | 3.56 | 7.60 |
| Weighted Avg. transgenic | 16 | 10.13 | 19.10 | 59.66 | 21.86 | 3.68 | 8.18 |
| % increase | | -18.3% | -23.9% | 13.7% | 11.7% | 3.2% | 7.7% |
| TTEST (family as e.u) | | 0.007 | 0.007 | 0.131 | 0.022 | 0.239 | 0.003 |
| TTEST (plant as e.u.) | | 3.3E-04 | 8.9E-05 | 2.9E-03 | 5.7E-06 | 1.5E-02 | 1.5E-08 |

**Supplementary Table 3.** Regeneration frequencies for different *GRF-GIF* combinations compared with empty vector in tetraploid wheat Kronos. The number of embryos used is indicated below each frequency. Regeneration frequencies were estimated as the number of calluses showing at least one regenerating shoot / total number of inoculated embryos. The blue "x" indicate the experiments included in the statistical analyses presented in the different figures and supplementary figures. All experiments in this Table used the regular 91 d protocol and *Agrobacterium* strain EHA105.

| | | GRF4- | Fig. | GRF4 & | Fig. | | | Fig. | GRF4- | | Fig. | GRF5 | GRF1 | GRF9 | Fig. |
|---|---|---|---|---|---|---|---|---|---|---|---|---|---|---|---|
| Exp. | pLC41 | GIF1 | 1D | GIF1 | 1E | GIF1 | GRF4 | 1F | GIF2 | GIF3 | 1G | | -GIF1 | | S1 |
| 1-a | 0.04<br>25 | 0.90<br>48 | x | | | | | | | | | | | | |
| 1-b | 0.08<br>25 | 0.96<br>25 | x | | | | | | | | | | | | |
| 2 | | 0.27<br>60 | | 0.06<br>32 | x | | | | | | | | | | |
| 3 | | 0.91<br>79 | | 0.77<br>83 | x | | | | | | | | | | |
| 3b[a] | | 0.60<br>47 | | 0.14<br>41 | x | | | | | | | | | | |
| 4 | 0.13<br>53 | 0.70<br>50 | x | 0.46<br>48 | x | 0.57<br>47 | | | | | | | | | |
| 6 | 0.20<br>20 | 0.65<br>20 | x | 0.50<br>20 | x | 0.35<br>20 | | | | | | | | | |
| 22 | 0.16<br>24 | 0.82<br>28 | x | | | 0.16<br>24 | 0.64<br>25 | x | | | | | | | |
| 25 | 0.00<br>15 | 0.70<br>20 | x | | | 0.40<br>15 | 0.06<br>15 | x | | | | | | | |
| 25b[a] | 0.00<br>15 | 0.35<br>17 | x | | | 0.00<br>15 | 0.00<br>15 | x | | | | | | | |
| 26 | 0.08<br>24 | 0.55<br>20 | x | | | 0.20<br>20 | 0.20<br>20 | x | | | | | | | |
| 26b[b] | 0.06<br>16 | 0.31<br>16 | x | | | 0.10<br>10 | 0.12<br>16 | x | | | | | | | |
| 12 | 0.00<br>10 | 0.50<br>10 | x | | | | | | 0.20<br>10 | 0.20<br>10 | x | | | | |
| 13 | 0.17<br>24 | 0.72<br>25 | x | | | | | | 0.50<br>24 | 0.46<br>24 | x | | | | |
| 24 | 0.10<br>10 | 0.50<br>10 | x | | | | | | 0.50<br>10 | 0.30<br>10 | x | . | 0.10<br>10 | 0.30<br>10 | x |
| 17 | 0.00<br>21 | 0.67<br>21 | x | | | | | | | | | 0.57<br>21 | 0.16<br>19 | 0.19<br>21 | x |
| 18 | 0.20<br>24 | 0.88<br>24 | x | | | | | | | | | 0.70<br>23 | 0.79<br>24 | 0.76<br>21 | x |
| 28 | 0.02<br>44 | 0.56<br>43 | x | | | | | | | | | 0.21<br>42 | 0.16<br>45 | 0.06<br>18 | x |

"-" indicates a fused protein or chimera, "&" indicates individual genes induced by separate promoters.

[a] No embryo dissection.

[b] No cytokinin.

**Supplementary Table 4.** Regeneration frequencies in plants transformed with the *Ubi::GRF4-GIF1* chimera or the empty vector pLC41. **A**) Tetraploid and hexaploid wheat commercial cultivars. **B**) Triticale breeding line UC3190. EHA105 and AGL1 are two different *Agrobacterium* strains (no differences were observed between the two strains). The number of embryos used for each genotype is indicated below the regeneration frequency.

**A. Wheat**

| Desert King (4x) | Exp1 EHA105 | Exp2 EHA105 | **Average** | **SE** |
|---|---|---|---|---|
| pLC41 | 0.05<br>20 | 0<br>50 | **0.025** | 0.025 |
| *Ubi::GRF4-GIF1* | 0.80<br>20 | 0.46<br>50 | **0.630** | 0.170 |

| Fielder (6x) | Exp1 UCD EHA105 | Exp2 UCD EHA105 | UCD **Average** | UCD **SE** |
|---|---|---|---|---|
| pLC41 | 0.05<br>49 | 0<br>10 | **0.025** | 0.025 |
| *Ubi::GRF4-GIF1* | 0.58<br>67 | 0.5<br>10 | **0.540** | 0.040 |

| Fielder (6x) | Exp1 JIC | Three Fielder experiments | |
|---|---|---|---|
| | AGL1 | **Average** | **SE** |
| pAGM8031 | 0.33<br>81 | 0.127 | 0.103 |
| *Ubi::GRF4-GIF1* | 0.775<br>40 | 0.618 | 0.082 |

| Cadenza (6x) | Exp1 AGL1 | Exp2 AGL1 | Exp3 EHA105 | **Average** | **SE** |
|---|---|---|---|---|---|
| pLC41 | 0<br>19 | 0<br>23 | 0<br>25 | **0.000** | 0.000 |
| *Ubi::GRF4-GIF1* | 0.20<br>12 | 0.17<br>24 | 0.20<br>25 | **0.190** | 0.010 |

| Hahn (6x) | Exp1<br>EHA105 | Exp2<br>EHA105 | Exp3<br>EHA105 | Exp4<br>AGL1 | **Average** | **SE** |
|---|---|---|---|---|---|---|
| pLC41 | 0<br>31 | 0<br>48 | 0<br>69 | 0<br>25 | **0.000** | 0.000 |
| *Ubi::GRF4-GIF1* | 0.03<br>37 | 0.04<br>50 | 0<br>51 | 0.28<br>25 | **0.088** | 0.087 |

#### Supplementary Table 4B. Triticale

| Triticale UC3190 (6x) | Exp9<br>EHA105 | Exp11<br>EHA105 | Exp15<br>EHA105 | **Average** | **SE** |
|---|---|---|---|---|---|
| pLC41 | 0<br>45 | 0<br>21 | 0<br>42 | **0.000** | 0.000 |
| *Ubi::GRF4-GIF1* | 0.05<br>45 | 0.13<br>22 | 0.14<br>43 | **0.107** | 0.028 |

**Supplementary Table 5.** Regeneration frequencies in wheat transformation with *Agrobacterium*

| Wheat methods | Explant | Average efficiency | Marker | Agro strain | Cultivars |
|---|---|---|---|---|---|
| This study without *GRF4-GIF1* | immature embryos | 8.3 / 2.5 %<br>12.7 % / 0.0 % | HPT | EHA105 | Kronos (4x) / 1 other (4x)<br>Fielder (6x) / 2 other (6x) |
| *GRF4-GIF1* | immature embryos | 65.1 / 63.0 %<br>61.8 / 13.9 % | HPT | EHA105 | Kronos (4x) / 1 other (4x)<br>Fielder (6x) / 2 other (6x) |
| Cheng et al., 1997 [2] | immature embryos | 2.2 % | NPT | C58 (ABI) | Bobwhite (6x) |
| Khanna HK, Daggard GE 2003 [3] | immature embryos | 3.9 % | PPT | LBA4404 | Veery5 (6x) |
| Wu et al., 2003 [4] | immature embryos | 9.5 / 4.5 % | PPT | AGL1 | Bobwhite / 3 other (6x) |
| Cheng et al., 2003 [5] | immature embryos | 1.1% | NPT II | C58 (ABI) | Bobwhite (6x) |
| Hu et al., 2003 [6] | immature embryos | 4.4 % | Glyphosate | C58 (ABI) | Bobwhite (6x) |
| Przetakiewicz et al., 2004 [7] | immature embryos | 12.6 / 2.3 % | NPT II | EHA101 / LBA4404 | Kontesa, Torka & Eta (6x) |
| Mitic et al., 2004 [8] | immature embryos | 0.6 % | PPT / HPT | AGL1 / LBA4404 | Vesna (6x) |
| Wu et al., 2008 [9] | immature embryos | 3.0 % | PPT | AGL1 | Ofanto (4x) |
| Risacher et al., 2009 [10] | immature seeds *in planta* | 5.0 % | NPT II | EHA105 | NB1 (6x) |
| He et al., 2010 [11] | immature embryos | 6.3 % | PPT | AGL1 | Stewart (4x) |
| Bińka et al., 2012 [12] | immature embryos | 3.4 % | NPT / PPT | EHA101/AGL1 | Kontesa, Torka (6x) |
| Hensel et al., 2017 [13] | immature embryos | 5 to 15 % [a] | HPT | AGL1 | Bobwhite (6x) |
| Hayta et al., 2019 [14] | immature embryos | 19 % | HPT | AGL1 | Fielder (6x) |
| **Proprietary Japan Tobacco [b]** | | | | | |
| Ishida et al., 2015 [c] [15] | immature embryos | 76.2 / 60.8 % | PPT / HPT | EHA101/EHA105 | Fielder (6x, PPT *vs* HPT) |
| Richardson et al., 2014 [16] | immature embryos | 40.9 / 12.1 %<br>50.8 / 26.0 % | PPT | AGL1 | Fielder / 9 other (6x)<br>Kronos / 1 other (4x) |
| Wang et al., 2017 [17] | immature embryos | 45.3 / 10.8 % | PPT | C58C1 | Fielder / 17 other (6x) |

[a] Only range provided

[b] At UCD, we purchased the JT license at and received training at their company. However, without the *GRF4-GIF1*, we have not been able to obtain the high regeneration efficiencies reported in Ishida et al. 2015 (likely because we use a wider range of embryo sizes collected from plant grown under different conditions)

[c] Report by the Japan Tobacco company in a non-peer reviewed journal

**Supplementary Table 6.** Regeneration frequencies in rice (*Oryza sativa*) cultivar Kitaake. Experiments 1 and 6 utilized the wheat-optimized vector pLC41 with or without the *Ubi::GRF4-GIF1* chimera. Experiments 2-5 used pCAMBIA1300, a vector frequently utilized in rice transformation, with or without the *Ubi::GRF4-GIF1* chimera. In each of these three experiments, calli generated from the same seed stock were inoculated with *Agrobacterium* containing the designated vector construct. In experiment 2, pCAMBIA1300-sgRNA refers to the pCAMBIA1300 vector carrying *Ubi::GRF4-GIF1* chimera plus a sgRNA targeted to gene *OsKitaake06g041700* encoding a TYROSYLPROTEIN SULFOTRANSFERASE (TPST). In experiments 3 and 4, pCAMBIA1300-gus refers to the control pCAMBIA1300-gus without the chimera. All experiments employed *Agrobacterium* strain EHA105. The number of calli used for each genotype is indicated below the regeneration frequency

| Rice Kitaake<br>No. calli inoc. | Exp1<br>n=85 | Exp2-4<br>n=100 x 3 | Exp5<br>n=50 | Exp6<br>n=50 | **Average** | **SE** |
|---|---|---|---|---|---|---|
| No *GRF4-GIF1* | 0.118<br>pLC41 | 0.235<br>pCAMBIA1300-gus | 0.22<br>pCambia1300 | 0.24<br>pLC41 | **0.2033** | 0.028 |
| *Ubi::GRF4-GIF1* | 0.353<br>pLC41 | 0.460<br>pCAMBIA1300-sgRNA | 0.44<br>pCambia1300 | 0.46<br>pLC41 | **0.4283** | 0.025 |

Paired *t*-test *GRF4-GIF1 vs.* control: $P < 0.0001$ (n = 4 experiments)

**Supplementary Table 7.** Regeneration frequencies in *Citrus*. Experiments 1 to 3 used a *GRF-GIF* chimera based on *Citrus* sequences whereas experiments 4 to 6 used a *GRF-GIF* chimera based on *Vitis* sequences. In addition, the last three experiments included a second *Vitis* construct with mutations in the miR396 binding site (*rGRF4-GIF1*) that precludes its cleavage. The number of epicotyls used for each genotype is indicated below the regeneration frequency.

| Carrizo | Exp1 | Exp2 | Exp3 | Exp4 | Exp5 | Exp6 | **Average** | **SE** |
|---|---|---|---|---|---|---|---|---|
| Empty vector | 0.04<br>45 | 0.00<br>38 | 0.12<br>56 | 0.02<br>65 | 0.09<br>32 | 0.02<br>40 | **0.05** | 0.02 |
| *Citrus GRF-GIF* | 0.15<br>45 | 0.39<br>41 | 0.16<br>69 | - | - | - | **0.21** | 0.09 |
| *Vitis GRF-GIF* | - | - | - | 0.07<br>59 | 0.25<br>31 | - | **0.16** | 0.09 |
| *Vitis* ***r****GRF4-GIF1* | - | - | - | 0.20<br>66 | 0.61<br>31 | 0.30<br>40 | **0.37** | 0.12 |

**Supplementary Table 8**. Comparisons of *GRF4-GIF1* with transformation technologies using different morphogenic genes.

| Technology | Ref. | Advantages | Disadvantages / limitations |
|---|---|---|---|
| GRF4-GIF1 | This one | 1. Publicly available for research<br>2. No developmental defects<br>3. Expands the range of genotypes that can be transformed<br>4. Rapid transformation protocol (60 days in wheat)<br>5. Robust regeneration efficiencies under broader set of protocols, including embryogenic and organogenic methods<br>6. Simple to implement and combine with gene editing<br>7. Efficient selection without selectable markers (wheat)<br>8. Tested in monocot and dicot species | 1. No tested yet in mature tissues in monocots<br>2. Transgene incorporated together with the *GRF4-GIF1* chimera [1].<br>3. Only tested in protocols that require in vitro tissue culture |
| *Bbm-Wus2*<br>(CORTEVA) | 18-20 | 1. High regeneration efficiencies in maize<br>2. Rapid transformation protocol (35 days maize)<br>3. Expanded range of maize germplasm that can be transformed<br>4. Works in mature tissues<br>5. Advanced vectors worked well in sorghum, Indica rice, and sugar cane | 1. Proprietary (but available for research)<br>2. Protocol optimized for maize. Use of specific maize promoters required to avoid regeneration problems and developmental defects<br>3. Tested only in monocots<br>4. If the BBM-WUS2 is not excised it induces developmental defects. Vectors with a CRE-LOX system are available<br>5. Only tested in methods that require in vitro tissue culture |
| *De novo* meristem induction<br>Fast-TrACC<br>*Wus2*/*ipt* | 21 | 1. Sidesteps the need for tissue culture<br>2. Co-delivery of developmental regulators and guide RNAs can generate edited shoots in plants constitutively expressing CAS9<br>3. It worked in *Benthamiana* soil grown plants | 1. Tested only in dicot plants (*Benthamiana*, tomato, potato, grape)<br>2. Fertile plants showed only in *Benthamiana*<br>3. Many edited plants show developmental defects and failed to produce seeds<br>4. Specific developmental regulators need to be defined in each new species<br>5. Needs transgenic plants previously transformed with CAS9 |

(1) This is not a problem for gene editing because both the CRISPR-CAS9 and the GRF4-GIF1 constructs are segregated out after editing. Although the presence of the *GRF4-GIF1* is not associated with developmental defects, the user can separate the transgene from the *GRF4-GIF1* chimera by using use a line previously transformed with the *GRF4-GIF1* without a selectable marker, and then retransforming the same line with the desired transgene. Since the transgene and the *GRF4-GIF1* construct are incorporated in different loci, they can be segregated apart.

### SUPPLEMENTARY METHODS

**Supplementary Method 1.** Vectors used in the transformation experiments.

***Wheat vectors.*** We performed all PCRs cloning with Phusion High-Fidelity DNA Polymerase (NEB). We extracted RNA extracted from spikes using the Spectrum Plant Total RNA Kit (Sigma-Aldrich), treated with RQ1 RNase-free DNase (Promega), and then synthesized the cDNA using SuperScript II Reverse Transcriptase (Invitrogen). To clone the coding region of wheat *GRF4* and *GIF1*, we performed PCRs using cDNA generated from Kronos spike. The sequence of the primers specific for *GRF4* (Fw-GRF4a/Rev-GRF4a) and *GIF1* (Fw-GIF1a/Rev-GIF1a) are indicated in Supplementary Table 1. We first cloned the PCR fragments in pDONR by a B/P gateway reaction and generated the *GRF4-GIF1* chimera by overlapping PCR.

In the first step, we amplified the *GRF4* and *GIF1* coding sequences with primers FW-GRF4a/Rev-GRF4b and Fw-GIF1b/Rev-GIF1b from the pDONR-GRF4 and pDONR-GIF1 clones. The primer Rev-GRF4b generates a 3' end that overlaps 12 nucleotides with the 5' end of Fw-GIF1b. Those 12 nucleotides generate a bridge of four alanine amino acids between GRF4 and GIF1. We gel-purified both PCR fragments and used them as template in a second PCR with the primers Fw-GRF4/Rev-GIF1b (Supplementary Table 1). We cloned the resulting product in pDONR. Next, we cloned the *GRF4*, the *GIF1* and the chimera *GRF4-GIF1* chimera the binary vector pLC41 by a L/R gateway reaction under the maize *UBIQUITIN* promoter. We verified the resulting vectors for the individual genes pLC41:*GRF4* and pLC41:*GIF1*, and for the chimera pLC41:*GRF4-GIF1* by restriction digestion, and transformed them by electroporation in *Agrobacterium* strain EHA105 and in a few experiments in strain AGL1 (Supplementary Table 4). Both strains were handled in the same way.

To develop the vector expressing both *GRF4* and *GRF1* under their own promoters (not fused, Ubi::*GRF4*-term and Ubi::*GIF1*-term), we amplified the complete Ubi::*GRF4*-term cassette by PCR using pLC41:*GRF4* as template with primers Fw_HindIII and Rev-HindIII (Supplementary Table 1). We cloned the PCR fragment in pGEMT-easy and then sub-cloned the Ubi::GRF4-term fragment into the *Hind*III site of pLC41:*GIF1*.

To generate the different wheat GRF-GIF chimeras, we obtained the coding sequences of *GRF1*, *GRF5*, *GRF9*, *GIF2* and *GIF3* by gene synthesis. Then, we generated the different chimeras (*GRF1-GIF1*, *GRF5-GIF1*, *GRF9-GIF1*, *GRF4-GIF2*, *GRF4-GIF3*) by overlapping PCR following the same strategy described to generate *GRF4-GIF1*. All the chimeras were cloned in pLC41 vector by L/R reaction. We verified all the vectors by restriction digestion and transformed by electroporation in *Agrobacterium* strain EHA105.

To develop the JD635-*GRF4-GIF1*-Cas9- gRNA-Gene *Q* vector, we amplified by PCR a cassette including the maize *UBIQUITIN* promoter, the *GRF4-GIF1* chimera and the Nos terminator (primers Fw_ZmUbi-AscI and Rev_NosTerm-AscI). The PCR product was gel-purified and cloned by In-fusion (Takara Bio USA, Inc.) into the *Asc*I site of the pYP25F binary vector, which contains a wheat codon optimized Cas9 (TaCas9) with two nuclear localization signals (NLS), and is a modified version of pDIRECT_25F (https://www.addgene.org/91143/) from Dr. Daniel Voytas group. We validated the vector sequence by Sanger sequencing. Next, we cloned a guide RNA construct targeting the coding region of Gene *Q* [22] by Golden Gate reaction into two *Aar*I sites of the vector and transformed it into chemical competent *E. coli* DH5α. We validated the JD635-*GRF4-GIF1*-Cas9-gRNA-Gene *Q* vector by Sanger sequencing and transformed by electroporation into *Agrobacterium* strain EHA105.

***Citrus and Vitis vectors.*** We generated the *Citrus* and *Vitis GRF-GIF* chimeras by gene synthesis using the *GRF* and *GIF* homologs highlighted in Supplementary Figure 1. We cloned the DNA fragments into pDONR by B/P gateway reaction. We cloned the *GRF-GIF* chimeras in the binary vector pGWB14 binary vector (L/R gateway reaction under viral 35S promoter) and transformed them by electroporation in *Agrobacterium* strain EHA105.

We generated a miR396-resistant version of *Vitis GRF-GIF* (*rGRF-GIF*) by overlapping PCR. Two PCR reactions were performed with primers Fw-GRF/rGRF-Rev and rGRF-Fw/Rev-GIF (Supplementary Table 1) using pGBW14-*vitis GRF-GIF* clone as template. The primers rGRF-Fw and rGRF-Rev overlap in 17 nucleotides, and introduce silent mutations in the miR396 target site (Supplementary Figure 6). We gel-purified both PCR fragments and used them as template in a second PCR with the primers Fw-GRF/Rev-GIF (Supplementary Table 1). We cloned the resulting product in pDONR by B/P gateway reaction. Next, we cloned the chimera *rGRF-GIF* in the binary vector pGWB14 by a L/R gateway reaction under the viral 35S promoter and transformed them by electroporation in *Agrobacterium* strain EHA105.

**Supplementary Method 2. Wheat transformation.** Wheat transformation followed previously published protocols [15]. Briefly, we grew the different wheat and triticale cultivars in a green house or a growth chamber under long-day photoperiod (16 h of 380 μM $m^{-2}$ $s^{-1}$ light, 26 °C day and 18 °C night). We harvested immature grains from spikes approximately 2 weeks after anthesis, and surface sterilized for 1 minute in 70 % ethanol followed by 10 minutes in 1.2% (v/v) sodium hypochlorite solution plus 5 μl tween. After surface sterilization, we washed the seeds three times with sterilized water and isolated immature embryos under stereoscopic microscope (embryo sizes 1.5 to 3.0 mm).

We centrifuged the isolated immature embryos in liquid medium and then inoculated with *Agrobacterium*. We transferred the embryos to co-cultivation medium with the scutellum-side up and incubated at 23 °C in the dark. After 2-3 days, we excised the embryo axis, and transferred them to callus induction medium without selection, where we incubated them at 25 °C in the dark. After 5 days, we transferred the embryos to selection medium with 30 mg/l of hygromycin and incubated them at 25 °C in the dark.

After 3 weeks, we transferred the calli to selection medium that contained 100 mg/l of hygromycin. After an additional 3 weeks, we transferred the proliferating tissue to regeneration medium containing 50 mg/l of hygromycin and incubated them at 25 °C under continuous light (30 μM $m^{-2}$ $s^{-1}$) for 2 weeks. We transferred the regenerated shoots into rooting medium contained 50 mg/l of hygromycin. Rooted plants were acclimated to soil by transferring them to a 1020 tray containing a 36 sheet inserts filled with Sunshine potting mix and covered with an 11 x 21 x 2 inch clear plastic dome for 10 days under 16 hour of 100 μM light and 26 °C. More recently we developed a shorter transformation protocol to generate *GRF4-GIF1* transgenic wheat plants that is summarized in Supplementary Figure 2.

Transformation at the John Innes Centre was performed as described by Hayta et al. (2019) [14].

**Supplementary Method 3. Rice transformation.** Rice transformation followed previously published protocols [23]. Briefly, we selected fresh rice seeds, de-husked them and surface sterilized them in a rotating flask containing 20 % (v/v) bleach for 30 min. Then, rinsed the seeds 3 times with sterile water. We placed about 25-50 seeds per plate on callus induction media (MSD, 1x Murashige and Skoog with vitamins medium containing 30 g/l sucrose, 2 mg/l 2,4-dichlorophenoxyacetic acid, 1.2% (w/v) agar, pH 5.6-5.8) without letting the embryo touch the media, wrapped plate with surgical tape end incubated under 16 h light/ 8 h dark at 28 ºC. After 10-14 d, we separated the callus from the rest of the germinating seed and transferred to fresh MSD agar plates for another 5 d before co-cultivation.

*Agrobacterium culture:* We prepared a glycerol freezer stock from a single bacterial colony isolated from a plate. We then inoculated 1 ml LB containing the appropriate antibiotics to maintain the *Agrobacterium* and the plasmid, and we incubated it overnight at 28 °C at 250 rpm. The following day we added 300 µl of the *Agrobacterium* culture to 20 ml TY (pH 5.5) containing the appropriate antibiotics and 200 µM acetosyringone. We incubated the culture 28 ºC for in a shaking incubator set at 250 rpm until the culture reached an $OD_{600}$ between 0.1 - 0.2 (approximately 2-4 h).

*Transformation and co-cultivation*: We placed the calli in *Agrobacterium* suspension for 30 min, and shook the suspension to ensure uniform access to the calli. After the shaking incubation, we dried the calli on sterile Whatman paper to remove excess bacterial suspension. We transferred the calli onto co-cultivation medium (MSD + S + AS, 1x Murashige and Skoog with vitamins medium containing 30 g/l sucrose, 5% sorbitol, 2mg/l 2,4-dichlorophenoxyacetic acid, 200 µM acetosyringone, 1.6 % (w/v) agar, pH 5.6-5.8) and incubated for 3 d in the dark at 22 ºC.

*Selection*: We transferred the co-cultivated calli to selection media (MSD + CH + PPM, 1x Murashige and Skoog with vitamins medium containing 30 g/l sucrose, 2 mg/l 2,4-dichlorophenoxyacetic acid, 400 mg/L carbenicillin, 200 mg/l timentin, 1ml/l Plant Preservative Mixture, 80 mg/L hygromycin, 1.2% agar, pH 5.6-5.8 ) and incubated the plates under continuous light at 28 ºC . We subcultured these calli onto fresh selection media every 8-9 d.

*Regeneration and Rooting:* After 4-5 weeks on selection media, resistant micro-calli of approximately 2- 5 mm wide started to appear. We picked these off the original callus and transferred them to Petri dishes with regeneration media (BN + S + CH, 1x Murashige and Skoog with vitamins medium containing 30 g/l sucrose, 5 % sorbitol, 3 mg/l BAP, 0.5 mg/l NAA, 400 mg/l carbenicillin, 200 mg/l timentin, 1 ml/l Plant Preservative Mixture, 50 mg/l hygromycin, 1.6 % (w/v) agar, pH 5.6-5.8), and incubated under continuous light at 28 ºC. We subculture these calli onto fresh regeneration media every 8-9 days. After 4-5 weeks, the calli that started to turn green, were transferred to regeneration media with reduced hygromycin (BN + S + CH, 1x Murashige and Skoog with vitamins medium containing 30 g/l sucrose, 5 %

sorbitol, 3 mg/l BAP, 0.5 mg/l NAA, 400 mg/l carbenicillin, 200 mg/l timentin, 1 ml/l Plant Preservative Mixture, 25 mg/l hygromycin, 1.6 % (w/v) agar, pH 5.6-5.8). When the shoot was properly developed, we transferred the regenerated plants to rooting media (MS + H, 1x Murashige and Skoog with vitamins medium containing 25 mg/l hygromycin, 1.2 % (w/v) agar, pH 5.6-5.) and incubated in 16 h light/ 8 h dark 28 ºC. When roots were well developed, we transferred the plants to soil.

**Supplementary Method 4. *Citrus* Transformation.** We placed seeds of Carrizo citrange rootstock in water to imbibe and then peel off the seed coats making sure not to remove the integument. We surface sterilized seeds in 0.6% (v/v) sodium hypochlorite solution plus 5-µl tween 20 by placing them in a 50 ml centrifuge tube and shaking at 100 rpms for 20 minutes. We rinse the seeds 3x in 150-200 ml of sterile distilled water. We placed seeds on agar solidified 1/2x Murashige and Skoog minimal organics medium (1/2x MSO) containing 15 g/l sucrose, 7 gm TC agar (pH 5.6-5.8), and push seeds slightly into the medium for more uniform germination. Incubate in the dark at 26 °C.

*Agrobacterium culture*: We prepared a glycerol freezer stock from a single bacterial colony isolated from a plate. We then used 40 µl of the stock to inoculate 20 ml of MGL medium (pH 7.0) containing the appropriate antibiotics to maintain the *Agrobacterium* and the plasmid, and we incubated overnight at 28 °C at 250 rpm. The following day, we removed 5 ml of the overnight growth and transferred it to 15 ml of TY medium (pH 5.5) containing the appropriate antibiotics and 200 µM acetosyringone. We incubated the culture overnight at 28 °C at 250 rpm and then diluted the overnight culture grown in TY medium to an $O.D_{600\ nm}$ of 0.1 to 0.2.

*Co-cultivation*: We collected 2-5 week old etiolated epicotyls and place in a petri dish containing 10 ml of the *Agrobacterium* solution prepared above (0.1-0.2 $OD_{600}$). We cut submerged epicotyls into 0.5 cm sections and soak for 10 min. We transferred the epicotyl sections onto co-cultivation medium consisting of Murashige and Skoog minimal organics medium (MSO) modified with 30 g/l sucrose 3.0 mg/l BAP, 0.1 mg/l NAA, and 200-µM acetosyringone pH 5.6-5.8. Incubate at 23 °C in the dark.

*Induction*: After 2-3 days, we transferred the epicotyl pieces to induction medium consisting of MSO modified with 30 g/l sucrose, 3.0 mg/l BAP, 0.1 mg/l NAA, 400 mg/l carbenicillin, 150 mg/l timentin and 100mg/l kanamycin sulfate, and incubated them in the dark. After 10 days, we subcultured the epicotyl sections to fresh medium of the same formulation and then subcultured them every 21 d. After the second 21-day cycle in the dark, we transferred cultures to light under a 30 µM light and a photoperiod of 16 h light 8 h dark. We continued to transfer every 21 d to fresh medium of the same media until organogenic shoot buds develop at the cut ends.

*Elongation*: Once shoots began to form, we transferred the developing shoots to elongation medium consisting of MSO modified with 30 g/l sucrose, 0.1mg/l BA, 400 mg/l carbenicillin, 150 mg/l timentin, and 100 mg/l kanamycin sulfate. We incubated as above and subcultured the cultures every 21 d as needed until shoots elongated.

*Rooting*: Once a shoot reached 2-4 cm in height, we harvested the shoots and transferred them to rooting medium consisting of MSO modified with 30 g/l sucrose, 5 mg/l NAA, 250 mg/l cefotaxime, and 100 mg/l kanamycin. After three to five days, we transferred shoots to MSO modified with 30 g/l sucrose, 0.0 mg/l NAA, 400 mg/l carbenicillin and 100 mg/l kanamycin. Shoots started rooting in 14 days.
